# Overexpression of Mdm36 reveals Num1 foci that mediate dynein-dependent microtubule sliding in budding yeast

**DOI:** 10.1101/2020.03.30.014621

**Authors:** Safia Omer, Katia Brock, John Beckford, Wei-Lih Lee

**Author notes:** University of Southern California School of Pharmacy, Clinical and Experimental Therapeutics Graduate Program (John Beckford).

## Abstract

Current model for spindle positioning requires attachment of the microtubule (MT) motor cytoplasmic dynein to the cell cortex, where it generates pulling force on astral MTs to effect spindle displacement. How dynein is anchored by cortical attachment machinery to generate large spindle-pulling forces remains unclear. Here, we show that cortical clustering of Num1, the yeast dynein attachment molecule, is limited by Mdm36. Overexpression of Mdm36 results in an overall enhancement of Num1 clustering but reveals a population of dim Num1 clusters that mediate dynein-anchoring at the cell cortex. Direct imaging shows that bud-localized, dim Num1 clusters containing only ∼6 copies of Num1 molecules mediate dynein-dependent spindle pulling via lateral MT sliding mechanism. Mutations affecting Num1 clustering interfere with mitochondrial tethering but not dynein-based spindle-pulling function of Num1. We propose that formation of small ensembles of attachment molecules is sufficient for dynein anchorage and cortical generation of large spindle-pulling force.

## INTRODUCTION

Orienting the mitotic spindle is paramount to controlling the outcome of asymmetric cell division, which is critical for determining daughter cell size, fate, and location. In most animals and fungi, pulling forces for orienting the spindle are generated by the microtubule motor cytoplasmic dynein, and are dependent on the anchoring of the dynein motor at the cell periphery (Bowman et al., 2006; Couwenbergs et al., 2007; Du and Macara, 2004; Heil-Chapdelaine et al., 2000; Kotak et al., 2012; Kotak et al., 2014; Nguyen-Ngoc et al., 2007; Sana et al., 2018). It has been proposed by recent work that cortical dynein-anchoring proteins might enhance spindle pulling function by forming clusters on the cell membrane (Okumura et al., 2018; Seldin et al., 2016); a favored hypothesis is that clustering of anchoring proteins contributes to the generation of large cooperative pulling forces by increasing the number of interacting motors per cortical MT contact site (Kiyomitsu, 2019; Okumura et al., 2018), analogous to how lipid microdomains on phagosomes facilitate motor clustering to achieve cooperative force generation of dynein during intracellular trafficking (Rai et al., 2016). However, despite identification of the domain required for clustering activity and punctate localization in both yeast and mammalian dynein-anchoring proteins (Harborth et al., 1999; Okumura et al., 2018; Tang et al., 2012), factors influencing cluster assembly, size, and distribution along the cortex are poorly understood. Additionally, whether cluster enhancement is correlated with increased dynein recruitment or higher cortical dynein activity remains unknown.

The budding yeast dynein-anchoring protein Num1 is a 313 kDa protein composed of a short N-terminal CC domain (aa 95-303), followed by a central TR domain (aa 592-1776) containing thirteen 64-residue tandem repeats, and a C-terminal PI(4,5)P2-binding PH domain (aa 2563-2683) (Greenberg et al., 2018). The CC domain is necessary and sufficient for cluster formation, in addition to being required for mediating an interaction with dynein and dynactin at the cell cortex (Tang et al., 2012). The CC domain also binds Mdm36, a 65 kDa protein implicated in promoting Num1 cluster formation and mitochondrial division (Hammermeister et al., 2010; Lackner et al., 2013). Although it is well-accepted that dynein exerts spindle pulling force at cortical Num1 sites, the abundance and heterogeneity of Num1 patches along the cell cortex (Heil-Chapdelaine et al., 2000; Omer et al., 2018; Schmit et al., 2018) has made it impossible to follow the effects of astral MT plus end interaction with individual cortical Num1 sites, a prerequisite for understanding how clustering might impact dynein force amplification. To our knowledge, contacts between astral MT plus end and individual Num1 foci have not been observed for MT sliding, the *in vivo* hallmark of dynein-mediated spindle pulling (Adames and Cooper, 2000; Yeh et al., 2000), hence the size of Num1 clusters required for this classic dynein-dependent microtubule-cortex interaction remains unknown. Additionally, recent work shows that organelles such as mitochondria and endoplasmic reticulum (ER) are involved in regulating Num1 cluster formation: a subset of cortical Num1 clusters appears to require mitochondria for their assembly (Kraft and Lackner, 2017), whereas another population requires the ER tethering proteins Scs2/Scs22 for their distribution throughout the cell cortex (Chao et al., 2014; Omer et al., 2018). The general hypothesis emerging from these studies is that distinct populations of Num1 clusters might exist at the cell periphery, but whether different pools of Num1 could be performing different Num1 functions – namely, dynein anchoring and mitochondrial tethering – remains a total mystery.

Here, we set out to characterize the role of Mdm36 in Num1 clustering and found that, in contrast to the prevailing notion for dynein-anchoring proteins, enhancing Num1 clustering unexpectedly reduces dynein recruitment to the cell cortex, but without affecting dynein function in spindle positioning. We report direct observation of MT sliding occurring upon astral MT plus end encountering a cortical Num1 cluster containing only a small number of Num1 molecules. The observed sliding events do not appear to require Mdm36 and mitochondria. Furthermore, mutations that interfere with Num1 clustering disrupt mitochondrial tethering activity but not dynein-based spindle-pulling activity of Num1, highlighting a more critical role for clustering in mitochondrial-anchoring rather than dynein-anchoring function.

## RESULTS

### Overexpression of Mdm36 dramatically enhances Num1 clustering

We first investigated how Num1 distribution is affected by Mdm36 at the cell cortex. As previously shown, deletion of Mdm36 resulted in smaller and dimmer Num1 patches (Lackner et al., 2013) (Fig. 1A), consistent with a clustering role for Mdm36. To determine whether Mdm36 level is limiting for Num1 clustering, we examined the effects of overexpressing Mdm36. We integrated the inducible *MET3* promoter at the 5’ end of the endogenous chromosomal locus of *MDM36* gene and assayed for Num1-GFP intensity and distribution. In Mdm36-overexpressing cells, hereafter referred to as Mdm36^OX^ cells, we observed a striking enhancement in the intensity of cortical Num1-GFP patches (Fig. 1A and Fig. S1A). The level of cytoplasmic Num1-GFP fluorescence was dramatically reduced compared to the levels in *mdm36*Δ null mutant and WT cells (Fig. 1A), suggesting that Mdm36-overexpression enhances the recruitment of free Num1 from the cytoplasm into cortical patches. We also observed that, similar to Mdm36-overexpression, overexpression of Num1 (using the same *MET3* promoter) resulted in an enhancement of Num1 patches (Fig. S1A). However, Num1 appeared to be more diffused, and the cortical patches were markedly more dispersed along the cortex in Num1-overexpressing cells compared to Mdm36^OX^ cells (Fig. S1B), indicating that Mdm36 is a clustering factor for Num1 at the cell membrane.

**Figure 1.**
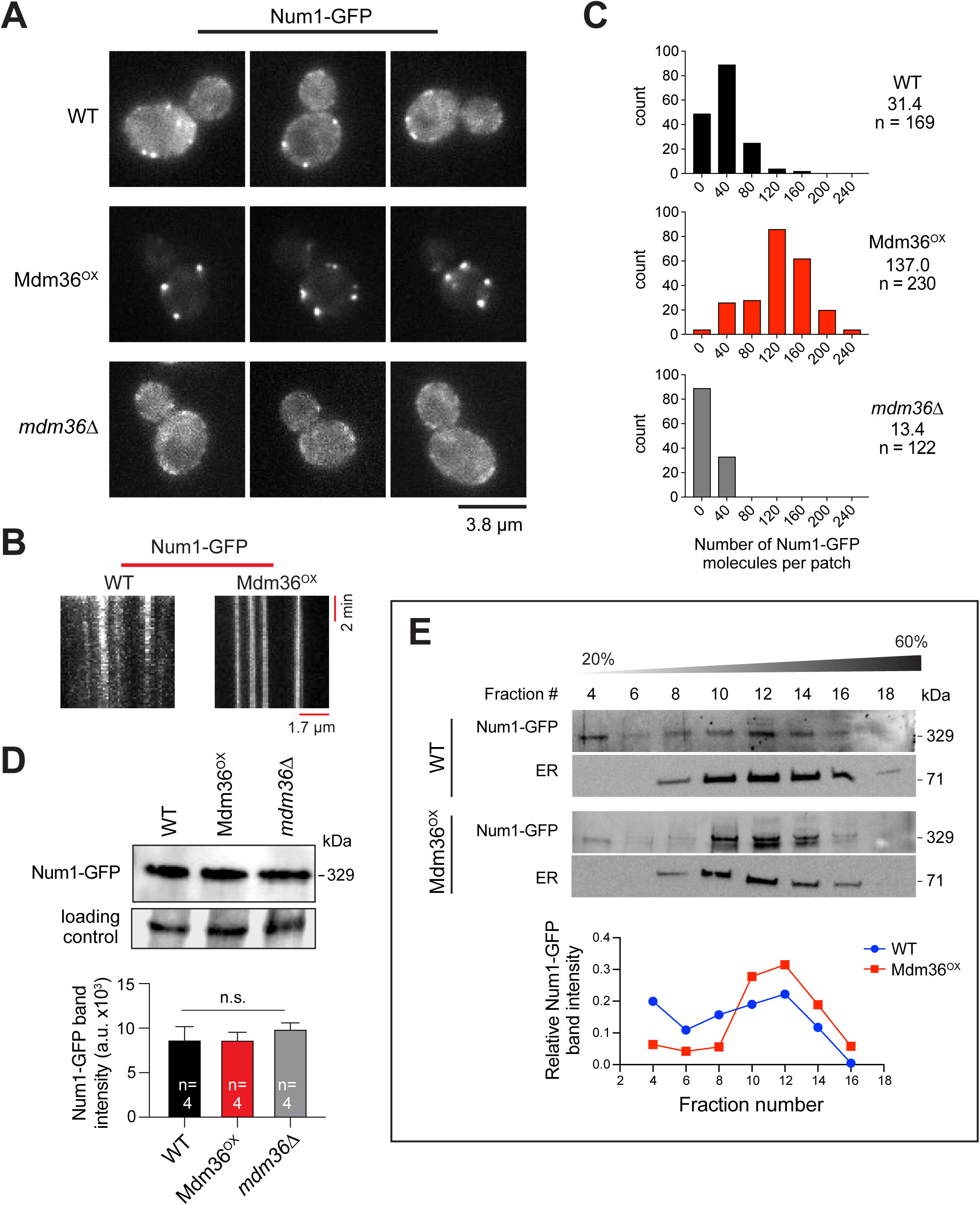
Mdm36 enhances Num1 clustering. (A) Representative single focal plane images of Num1-GFP in WT, Mdm36^OX^, and *mdm36*Δ cells acquired using the same camera settings. (B) Kymograph of Num1-GFP foci in WT and Mdm36^OX^ cells. (C) Histogram of protein copy number per cortical patch for Num1-GFP. Median copy number shown for each strain. (D) Western blot showing Num1-GFP protein level in whole cell lysate of indicated strains. Plot depicts mean intensity of Num1-GFP band for four loading replicates. Error bar represents standard error of the mean (SEM). n.s., not statistically significant by one-way ANOVA test. (E) Sucrose gradient sedimentation analysis of Num1-GFP in WT and Mdm36^OX^ strains. Whole cell lysate was loaded onto a 20-60% sucrose gradient, sedimented, and analyzed by western blot using anti-GFP (for Num1-GFP) and anti-Sac1 (for ER) antibodies. *Bottom*, intensity of Num1-GFP band plotted against fraction number.

Kymograph analysis showed that Num1-GFP patches in Mdm36^OX^ cells were stationary, similar to Num1 patches in WT cells (Fig. 1B), and were remarkably stable, as some were observed persisting for more than two cell division cycles (Video 1). We used a quantitative ratiometric approach to determine the number of fluorescent Num1-GFP molecules at individual cortical patches in Mdm36^OX^ cells. Anaphase Cse4-GFP spots were used as a standard for the molecule counting assay (Verdaasdonk et al., 2014). In WT cells, we found 31 copies of Num1-GFP per cortical patch (Fig. 1C), a level higher than what we had previously reported (Tang et al., 2009) given the revised protein copy number of Cse4-GFP (Lawrimore et al., 2011; Verdaasdonk et al., 2014). The intensities of Num1-GFP patches in Mdm36^OX^ cells yielded 137 copies per cortical patch (Fig. 1C), a 4.4-fold increase from WT cells. In contrast, in *mdm36*Δ cells, we found 13 copies of Num1-GFP per cortical patch (Fig. 1C), a 2.3-fold decrease from WT cells. Immunoblot analysis of total cell lysates showed that Num1 levels in Mdm36^OX^ and *mdm36*Δ cells were similar to that in WT (Fig. 1D), indicating that the observed difference in the copy number could not be attributed to changes in the expression levels or the stability of Num1 protein. Thus, our data indicate that the copy number of Num1 per cortical patch is limited by the levels of Mdm36 in the cell.

Similar to Num1 patches in WT cells (Omer et al., 2018), Num1-GFP patches in Mdm36^OX^ cells redistributed to the polarized ends of the cell (the distal bud tip and the mother cell apex) upon deletion of the ER tethering proteins, Scs2 and Scs22 (Fig. S1C). To further assay for association with ER, we analyzed sedimentation profiles of Num1 in sucrose density gradients. The ER-localized PI phosphatase Sac1 was used as a marker for ER. We found that Num1 in Mdm36^OX^ lysate co-fractionated with ER, more so than Num1 in WT lysate (Fig. 1E; compare Num1-GFP levels in fractions 10, 12, and 14 to fractions 4, 6 and 8). These data indicate that Num1’s association with ER is enhanced by Mdm36-overexpression.

### Dynein function is intact in Mdm36^OX^ cells despite asymmetric distribution of Num1

We next wondered whether the observed change in Num1 clustering in Mdm36^OX^ cells would affect dynein pathway function. The prevailing model for dynein function proposes that dynein is recruited from the dynamic plus ends of astral MTs to cortical patches of Num1 in the bud; once anchored, dynein uses its minus end-directed motor activity to pull the attached spindle into the bud neck (Lee et al., 2005; Lee et al., 2003; Sheeman et al., 2003). In Mdm36^OX^ cells, however, we noticed that the enhanced Num1 patches were observed mostly in the mother cell compartment (Fig. 1A). The bud often exhibited no visible patches (Fig. 2A; compare intensity plot along the bud cortex for Mdm36^OX^ versus WT) or only a few dim patches during the entire budding process (Fig. 2B). Although the reason for the apparent asymmetry in Num1 distribution is unclear at this point, this phenomenon raises the possibility that dynein pathway function might be defective in the Mdm36^OX^ background.

**Figure 2.**
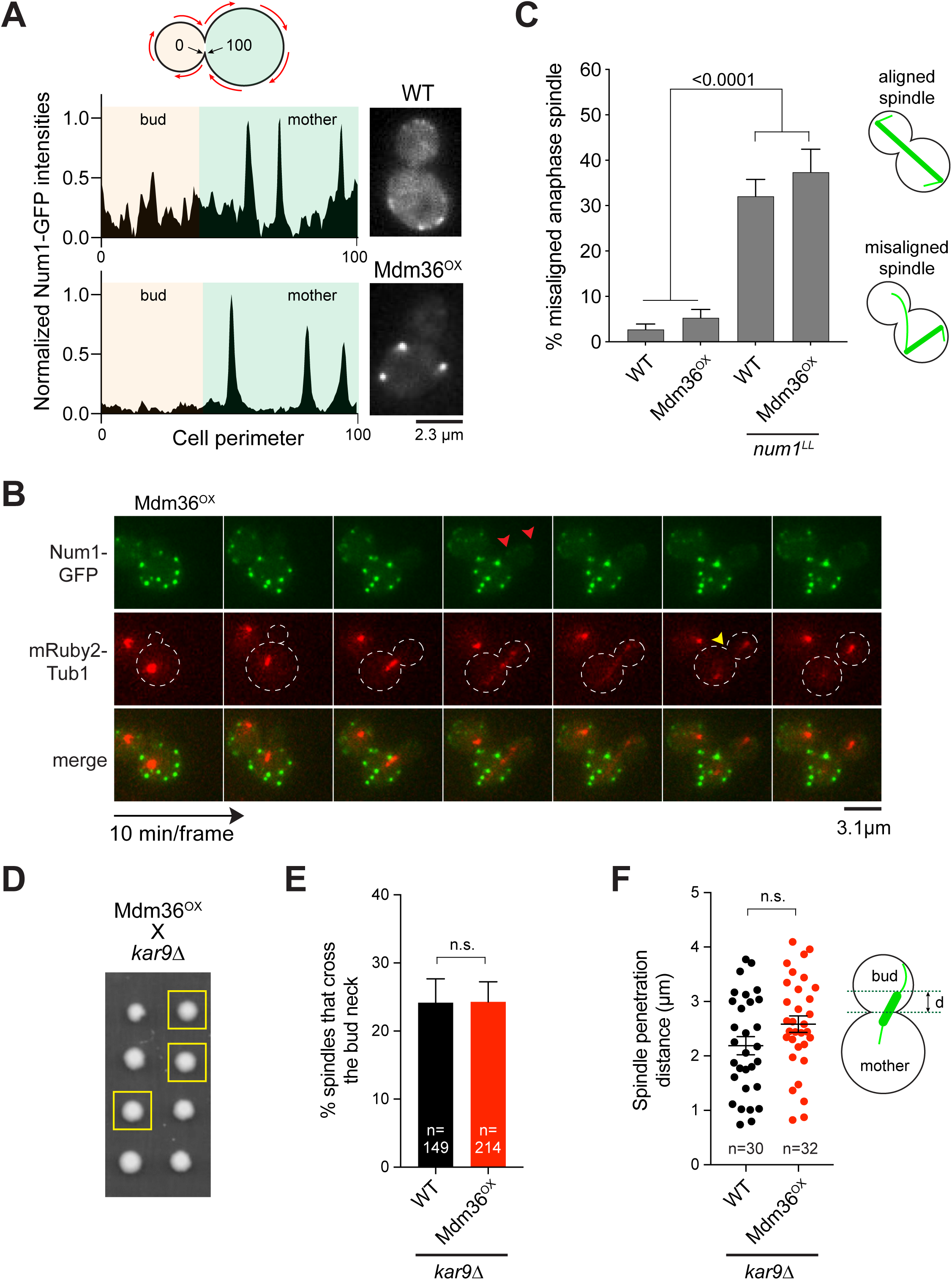
Dynein pathway function in Mdm36^OX^ cells. (A) Normalized intensity profile along the cell cortex (left) and single focal plane image (right) of a representative WT and Mdm36^OX^ cell expressing Num1-GFP. (B) Asymmetry of Num1 localization between mother and daughter cell in Mdm36^OX^ background during the budding cycle. *Red arrowheads*, dim Num1 patches in the bud. *Yellow arrowhead*, end of mitosis as indicated by spindle disassembly. (C) Percentage of anaphase spindles with a misoriented phenotype for indicated strains (91 ≤ n ≤ 514 spindles per strain). Error bar represents standard error of proportion (SEP). *P*-value by one-way ANOVA test. n.s., not statistically significant. (D) Representative tetrad progeny of a cross between Mdm36^OX^ (*MET3p:MDM36*) and *kar9*Δ. Yellow boxes indicate *MET3p:MDM36 kar9*Δ progeny as determined by marker analysis. (E) Percentage of HU-arrested spindles that crossed the bud neck during a 10-min movie in WT and Mdm36^OX^ cells in the *kar9*Δ background. (F) Spindle penetration distance in HU-arrested cells of indicated strains. Distance of spindle penetration (d) is defined as the farthest distance traveled by a spindle pole moving across the bud neck during a 10-min movie. n.s., not statistically significant.

We first assessed dynein pathway function using a simple spindle orientation assay, observing anaphase spindle position at a single time-point in a population of asynchronous cells. Remarkably, Mdm36^OX^ strain exhibited only 5.2% of cells with a misoriented anaphase spindle phenotype, quantitatively similar to that observed for a WT strain (2.7%; Fig. 2C), indicating that dynein pathway function is not defective. In contrast, Mdm36^OX^ cells expressing Num1^LL^, which harbors two point mutations that abolish the Num1-dynein interaction (Tang et al., 2012), exhibited a high level of misoriented anaphase spindle phenotype (37.4%; Fig. 2C), confirming that Num1-dynein interaction is required for proper spindle orientation in the Mdm36^OX^ background.

We further evaluated dynein function by assaying for synthetic growth defects with *kar9*Δ and *cin8*Δ. Budding yeast harboring *KAR9* or *CIN8* deletion requires the dynein pathway for normal growth (Geiser et al., 1997; Gerson-Gurwitz et al., 2009; Miller and Rose, 1998). Tetrad dissection analysis showed that *MDM36*^*OX*^ *kar9*Δ and *MDM36*^*OX*^ *cin8*Δ progeny formed viable haploid colonies (Table I), exhibiting no growth defects when compared with *kar9*Δ and *cin8*Δ single mutants (Fig. 2D), consistent with the dynein pathway being functional in the Mdm36^OX^ background.

**Table I.**
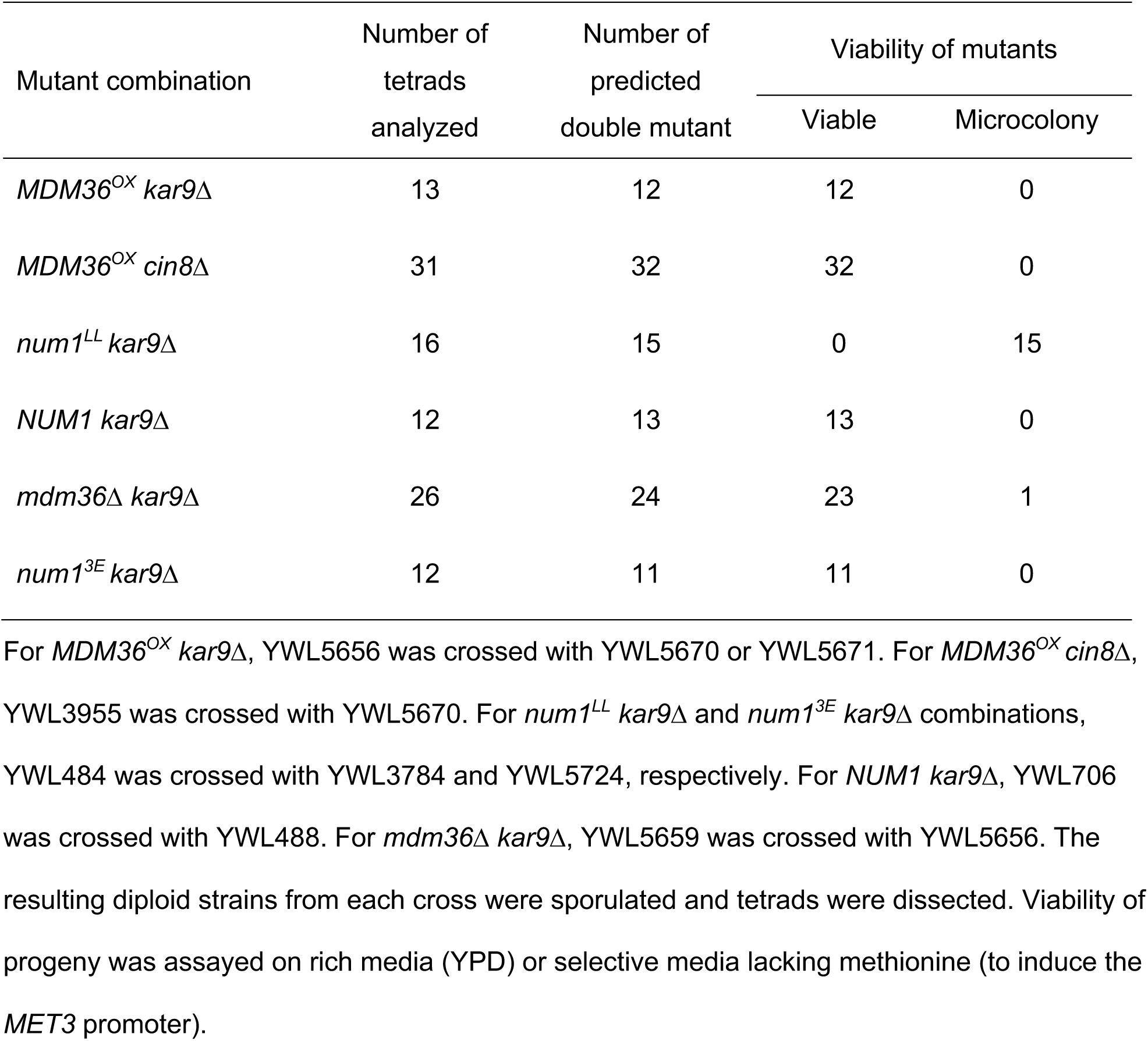
Viability of mutants in combination with *kar9*Δ

We next quantitated dynein function using a spindle crossing assay, scoring for preanaphase spindle movements through the bud neck in a *kar9*Δ background. We found that Mdm36^OX^ *kar9*Δ cells exhibited the same frequency of spindle traversing the bud neck when compared with *kar9*Δ cells (24.3% versus 24.2%; Fig. 2E). Moreover, in cells where the spindle was able to penetrate the bud neck, it moved for a similar distance compared with *kar9*Δ cells (Fig. 2F), consistent with an intact dynein pathway function. These results are surprising given the observed asymmetry in Num1 distribution in Mdm36^OX^ cells.

### Dynein and dynactin localization are altered in Mdm36^OX^ cells

We asked whether the observed change in Num1 localization in Mdm36^OX^ cells affects dynein and dynactin targeting to the astral MT plus ends and the cell cortex. Strikingly, compared to WT cells, the frequency of observing cortical Dyn1-3GFP (dynein heavy chain) and Jnm1-3mCherry (dynactin p50^dynamitin^ subunit) foci was significantly reduced in the Mdm36^OX^ background (Fig. 3A). Furthermore, the mean fluorescence intensity of Dyn1-3GFP and Jnm1-3mCherry foci at the MT plus ends was significantly enhanced in Mdm36^OX^ relative to WT (Fig. 3B and 3C). These data indicate that the delivery of dynein and dynactin from the MT plus ends to the cell cortex via the offloading mechanism is reduced even though Num1 clustering is enhanced in Mdm36^OX^ cells. In agreement with the observed reduction in dynein-offloading, the mean intensity of the cortical Dyn1-GFP foci in Mdm36^OX^ cells was significantly decreased (Fig. 3B). The mean intensity of the cortical Jnm1-3mCherry foci appeared to decrease slightly, however since the cytoplasmic background fluorescence for Jnm1-3mCherry was high, the observed change was not significantly different from WT cells (Fig. 3C; p = 0.368). We next asked whether cortical dynein colocalizes with Num1 patches in the Mdm36^OX^ background, as would be expected if Num1 anchors dynein to the cell cortex. We scored for cortical Dyn1 by analyzing time-lapse two-color images of Dyn1-3GFP and Num1-mRuby3. To be considered cortical, the Dyn1-3GFP foci had to remain stationary at the cell cortex for ≥ 3 min. We observed that all cortical Dyn1-3GFP foci colocalized with Num1-mRuby3; however, to our surprise, the intensity of the Num1-mRuby3 patches that possessed Dyn1-3GFP was strikingly lower than those lacking Dyn1-3GFP (by ∼2.6-fold, Fig. 3D; n = 338 patches). Colocalization of Dyn1-3GFP with a bright Num1-mRuby3 cluster was rarely observed, suggesting that Mdm36^OX^-mediated Num1 clustering negatively regulates cortical dynein targeting. These analyses uncovered a compromised dynein targeting in the Mdm36^OX^ cells, albeit without causing a spindle misorientation phenotype.

**Figure 3.**
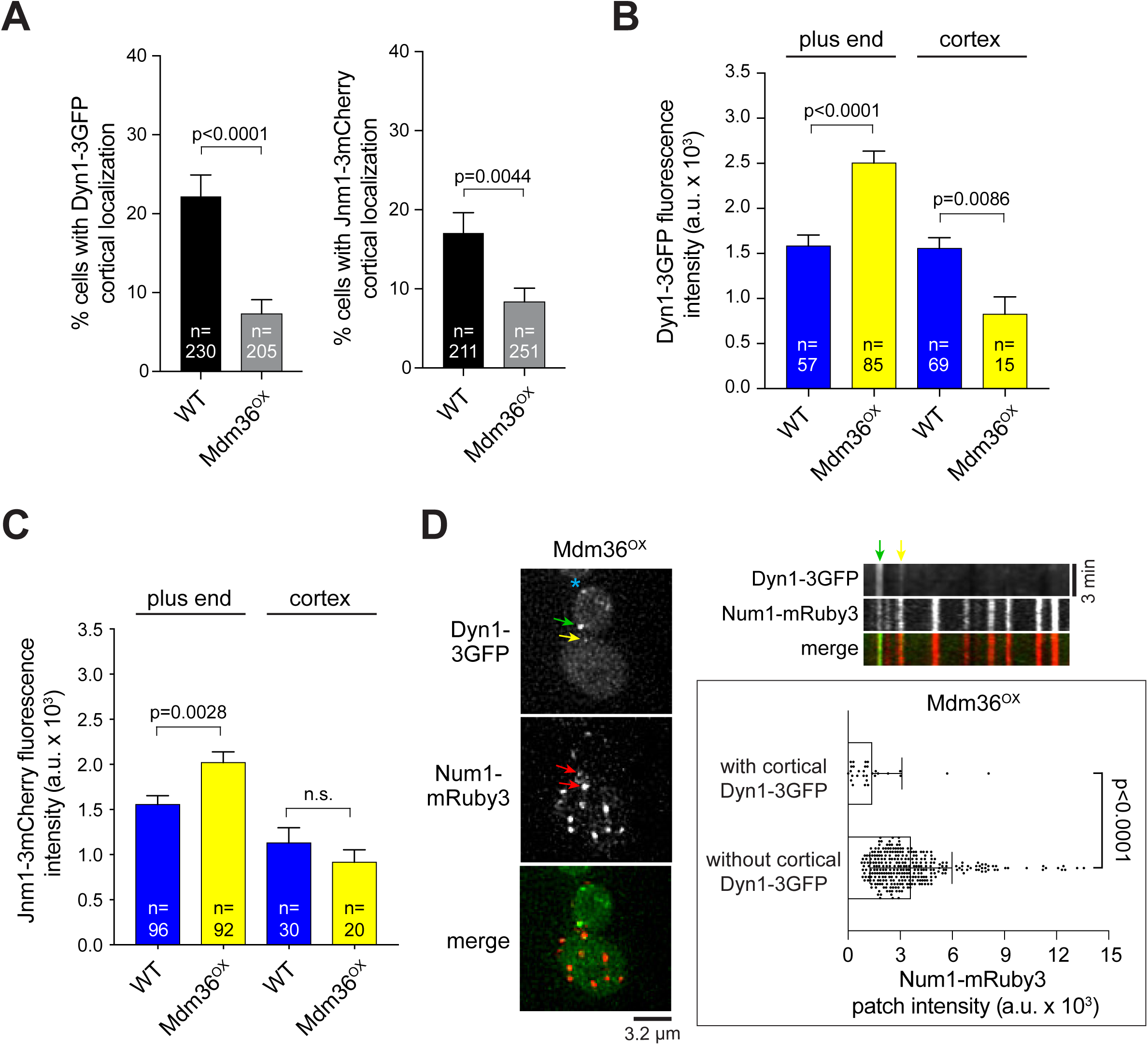
Defects in dynein and dynactin localization in Mdm36^OX^ cells. (A) Frequency of observing cortical Dyn1-3GFP and Jnm1-3mCherry patches. Error bar represents SEP. *P*-value by unpaired *t* test. (B and C) Dyn1-3GFP and Jnm1-3mCherry mean fluorescence intensity at the plus end and cell cortex. Error bar represents SEM. *P*-value by unpaired *t* test. n.s., not statistically significant. (D) *Left*, deconvolved wide-field images of a representative Mdm36^OX^ cell expressing Dyn1-3GFP and Num1-mRuby3. Green and yellow arrows mark stationary cortical Dyn1-3GFP foci (see kymograph, *top right*). Asterisk marks motile plus end Dyn1-3GFP focus (based on time-lapse sequence). Red arrows mark Num1-mRuby3 clusters possessing Dyn1-3GFP signal. *Top right*, kymograph of Dyn1-3GFP and Num1-mRuby3 foci from the same cell shown on the left. *Bottom right*, mean intensity of Num1-mRuby3 foci that possess or lack Dyn1-3GFP signal in Mdm36^OX^ cells (n = 338). Error bar represents standard deviation (SD). *P*-value by unpaired *t* test.

### Direct observation of MT sliding initiated by dim Num1 clusters

We next sought to determine how dynein mediates spindle positioning in Mdm36^OX^ cells. In WT cells, cortical dynein pulls the spindle into the bud cell compartment via lateral MT sliding mechanism (Adames and Cooper, 2000). More recently, however, several studies showed that cortical dynein can also pull on the astral MT via end-on MT capture-shrinkage mechanism (Laan et al., 2012; Omer et al., 2018). To assess the mechanism in Mdm36^OX^ cells, we monitored dynein-dependent astral MT interaction with the bud cortex using a spindle correction assay, scoring for MT behavior during anaphase spindle re-alignment from a misoriented position. In control *kar9*Δ cells, spindle correction was primarily mediated by MT sliding along the bud cortex (97.2 %, n = 36 events; Fig. 4A), as previously reported (Omer et al., 2018; Yeh et al., 2000). Remarkably, in *kar9*Δ Mdm36^OX^ cells, the mechanism of spindle correction was unaffected, despite the apparent asymmetry in Num1 distribution between the mother and bud compartments (Fig. 4A; 46 out of 52 misaligned spindles were corrected by MT sliding mechanism). Disrupting dynein-anchoring in Mdm36^OX^ cells using the *num1*^*LL*^ allele abolished lateral MT sliding and prevented spindle correction (0 out of 50 spindles were corrected in *num1*^*LL*^ Mdm36^OX^ cells; Fig. 4A). These results indicate that dim Num1 patches in the bud of Mdm36^OX^ cells are sufficient for dynein attachment and generation of spindle pulling force. As control, we found that spindle correction was also mediated by the MT sliding mechanism in *kar9Δ mdm36*Δ cells (18 out of 22 events, 82%), suggesting that the observed phenotype in *kar9*Δ Mdm36^OX^ cells was not due to an artificial effect of Mdm36 overexpression.

**Figure 4.**
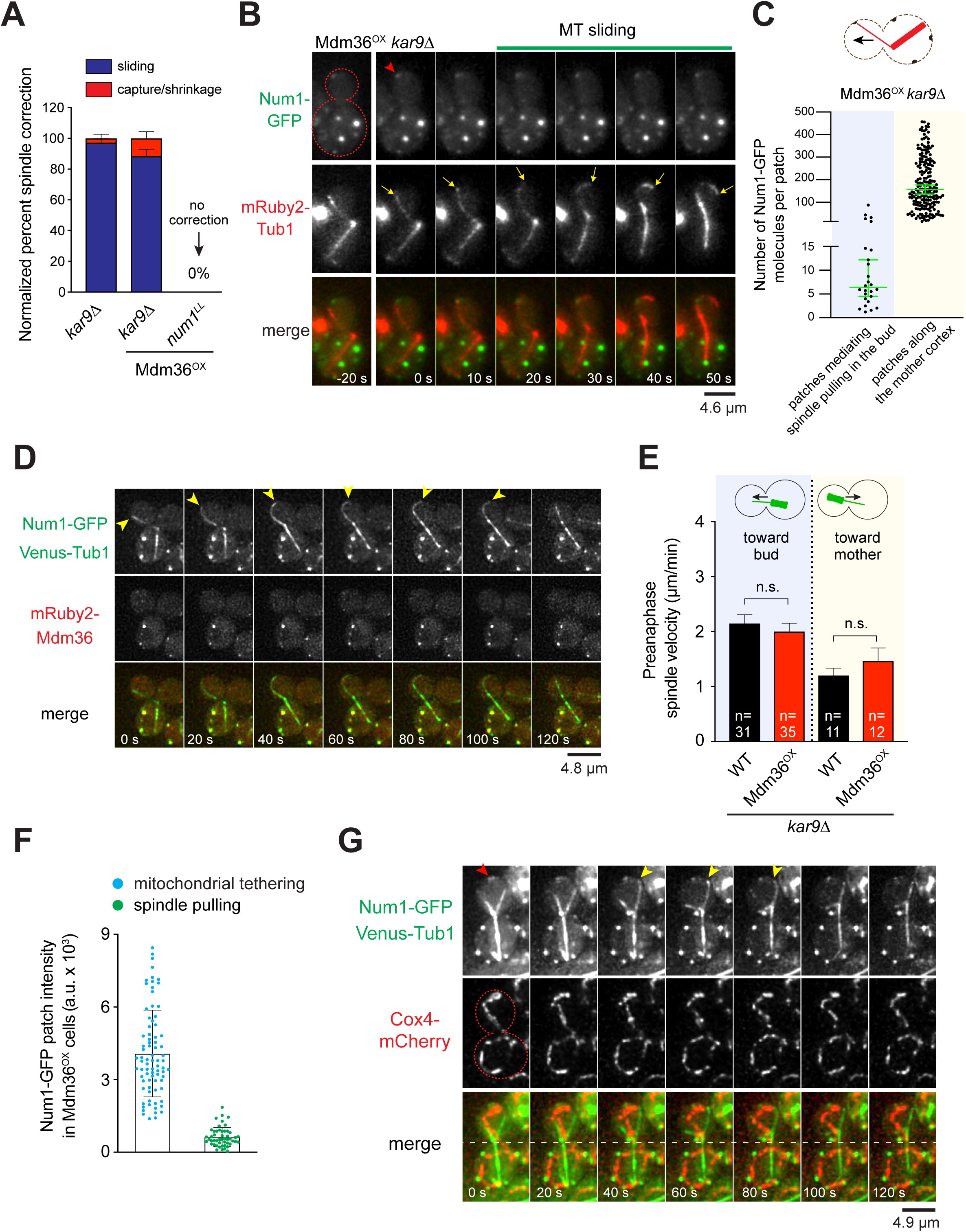
Dim Num1 patches mediate spindle pulling via MT sliding. (A) Quantification of spindle correction mechanisms for indicated strains (22 ≤ n ≤ 52 events per strain). Error bar represents SEP. (B) Time-lapse sequence showing spindle correction via MT sliding in a Mdm36^OX^ *kar9*Δ cell expressing Num1-GFP and mRuby2-Tub1. Red arrowhead marks initiation of MT sliding at a dim cortical Num1 cluster. Yellow arrows mark the position of the astral MT plus end. (C) Copy number of Num1-GFP for individual foci found along the mother cell cortex versus those found at the bud cortex mediating spindle pulling during a spindle correction assay in Mdm36^OX^ *kar9*Δ cells (n = 201 and 26, respectively). Median ± 95% confidence interval is indicated. (D) Deconvolved two-color image sequence showing spindle correction via MT sliding in a *NUM1-GFP VENUS-TUB1 kar9*Δ cell overexpressing mRuby2-tagged Mdm36. Arrowheads mark the position of the astral MT plus end. (E) Preanaphase spindle velocity toward the bud or mother cell observed during a 7-min movie. Error bar represents SEM. n.s., not statistically significant by unpaired *t* test. (F) Mean fluorescence intensity of Num1-GFP patches seen mediating spindle pulling or mitochondrial tethering in Mdm36^OX^ cells. Patches mediating spindle pulling were observed in *kar9*Δ background. Error bar represents SD. (G) Deconvolved two-color image sequence showing spindle pulling event in a Mdm36^OX^ *kar9*Δ cell expressing Num1-GFP, Venus-Tub1, and Cox4-mCherry. Yellow arrowheads indicate spindle pulling by MT sliding (frames 40 s to 80 s) in a cortical region lacking mitochondria. Red arrowhead marks Num1 foci involved in mitochondrial tethering.

To confirm this idea, we captured two-color time-lapse movies of *kar9*Δ Mdm36^OX^ cells that also expressed Num1-GFP. In these movies, we observed that MT sliding occurred upon the plus end encountering a dim Num1-GFP focus at the bud cortex (Fig. 4B; Video 2). The astral MT slid over the Num1-GFP focus, pulling the minus-end-attached spindle into the bud, causing spindle correction. We also sometimes observed MT sliding occurring over a cortical region without any apparently visible Num1-GFP focus (Video 3), suggesting that the signals from the dynein-anchoring Num1 foci were at or below our detection limit. Co-imaging with Cse4-GFP cells followed by ratiometric comparison of fluorescence intensity revealed that the Num1 foci in the bud that mediated spindle correction contained a significantly smaller copy number of Num1 molecules compared to the Num1 patches found along the mother cell cortex. The median copy number for the Num1 foci mediating spindle correction was 6 (n = 26), whereas the median copy number for the Num1 patches in the mother cell was 158 (n = 201; Fig. 4C). Thus, Num1 does not need to assemble into a large focal patch or structure to enable dynein to generate cortical spindle pulling forces. Additionally, further localization analysis revealed that, during spindle pulling events in *kar9*Δ Mdm36^OX^ cells, Mdm36 appeared to be absent at the site where MT sliding was initiated in the bud (Fig. 4D; 17 out of 24 events occurring without Mdm36). These results implicate the existence of a morphologically and functionally distinct population of Num1 foci mediating dynein-based spindle-pulling activity along the bud cell cortex.

### Num1 clustering enhances mitochondrial tethering but not dynein-mediated spindle-pulling function

To further examine whether patch enhancement has any effect on cortical dynein activity, we quantified spindle movements during spindle oscillation across the bud neck in HU-arrested *kar9*Δ Mdm36^OX^ cells. We considered the possibility that the bright Num1 patches in the mother compartment of *kar9*Δ Mdm36^OX^ cells might enable a stronger spindle-pulling force by dynein, potentially retarding the spindle traversing the neck from the mother to the daughter or, conversely, enhancing its movement from the daughter to the mother. However, we found that spindle oscillation across the bud neck was unaffected in *kar9*Δ Mdm36^OX^ cells compared with *kar9*Δ cells (Fig. 4E). Notably, the preanaphase spindle velocity for retrograde movements from the bud to the mother in *kar9*Δ Mdm36^OX^ cells was indistinguishable from that in *kar9*Δ cells (Fig. 4E), indicating that the bright Num1 patches observed in the mother cortex of *kar9*Δ Mdm36^OX^ did not correlate with an enhancement of cortical dynein activity.

To test whether patch enhancement correlates with mitochondrial tethering activity, we imaged Mdm36^OX^ cells expressing Num1-GFP and mitochondria-targeted Cox4-mCherry. We observed that Mdm36^OX^ cells displayed a branched and tubular mitochondrial network that localizes close to the cell periphery as observed in WT cells (Video 4; Fig. S2A); however, the mitochondrial network in Mdm36^OX^ cells exhibited an enhanced tethering phenotype: 88.9% of Mdm36^OX^ cells versus 55.0% of WT cells showed >3 persistent mitochondrial tether points over the course of a 3-min video (p < 0.0003, n ≥ 40 cells). Additionally, full 3D stacks showed that every bright Num1 patches at the cell cortex of Mdm36^OX^ cells was associated with mitochondria (Videos 4 and 5). Using the same camera settings and imaging conditions to capture mitochondrial tethering events in *MDM36*^*OX*^ *NUM1-GFP COX4-mCherry* cells and spindle correction events in *MDM36*^*OX*^ *NUM1-GFP mRuby2-TUB1 kar9*Δ cells, we found a striking relationship between the intensity of the Num1 patch and the function of the patch in the cell. When the intensity of 130 Num1 patches was plotted against the activity that they performed in the cell, we observed a strong association between the bright Num1 patches and mitochondrial tethering function, and between the dim Num1 patches and spindle-pulling function (Fig. 4F). On average, the Num1 patches mediating mitochondrial tethering were 6.4-fold brighter than the Num1 patches mediating spindle correction function. Furthermore, during spindle pulling events in *kar9*Δ Mdm36^OX^ cells, we observed that MT sliding was initiated at a cortical site devoid of mitochondria (Fig. 4G). These results, combined with the observation that patch enhancement in the mother compartment has no effect on the spindle velocity in either anterograde or retrograde direction (Fig. 4E), reveals a correlation between the size of the Num1 patch with mitochondrial tethering activity but not with dynein-based spindle-pulling activity.

### 3E mutations disrupt Num1 clustering but not cortical dynein activity

Notably, recent studies (Kraft and Lackner, 2017; Ping et al., 2016; Schmit et al., 2018) proposed that the assembly of functional Num1 patches in the bud is dependent on an interaction between Num1 and mitochondria. In light of our data suggesting differential functions of Num1 patches, we wondered if the interaction with mitochondria is required for the formation of the two morphologically and functionally distinct patches observed in Mdm36^OX^ cells. If mitochondria are indeed required for the formation of both types of patches, Mdm36^OX^ cells harboring disrupted Num1-mitochondria interaction should be defective in mitochondrial-tethering and spindle-pulling functions. To test this, we used a well-characterized mitochondria-binding mutant, Num1^K121E+R262E+R265E^ (hereafter referred to as Num1^3E^; Fig. 5A), which harbors three point mutations that disrupt the Num1-mitochondria interaction but do not interfere with the Num1-Mdm36 interaction (Ping et al., 2016). The 3E mutations were originally identified in a mutagenesis screen for mutations in the CC domain of Num1 that abolish its affinity for liposomes mimicking the phospholipid composition of the mitochondrial outer membrane (Ping et al., 2016), however how they affect Num1 cluster formation and dynein pathway function have not been characterized.

**Figure 5.**
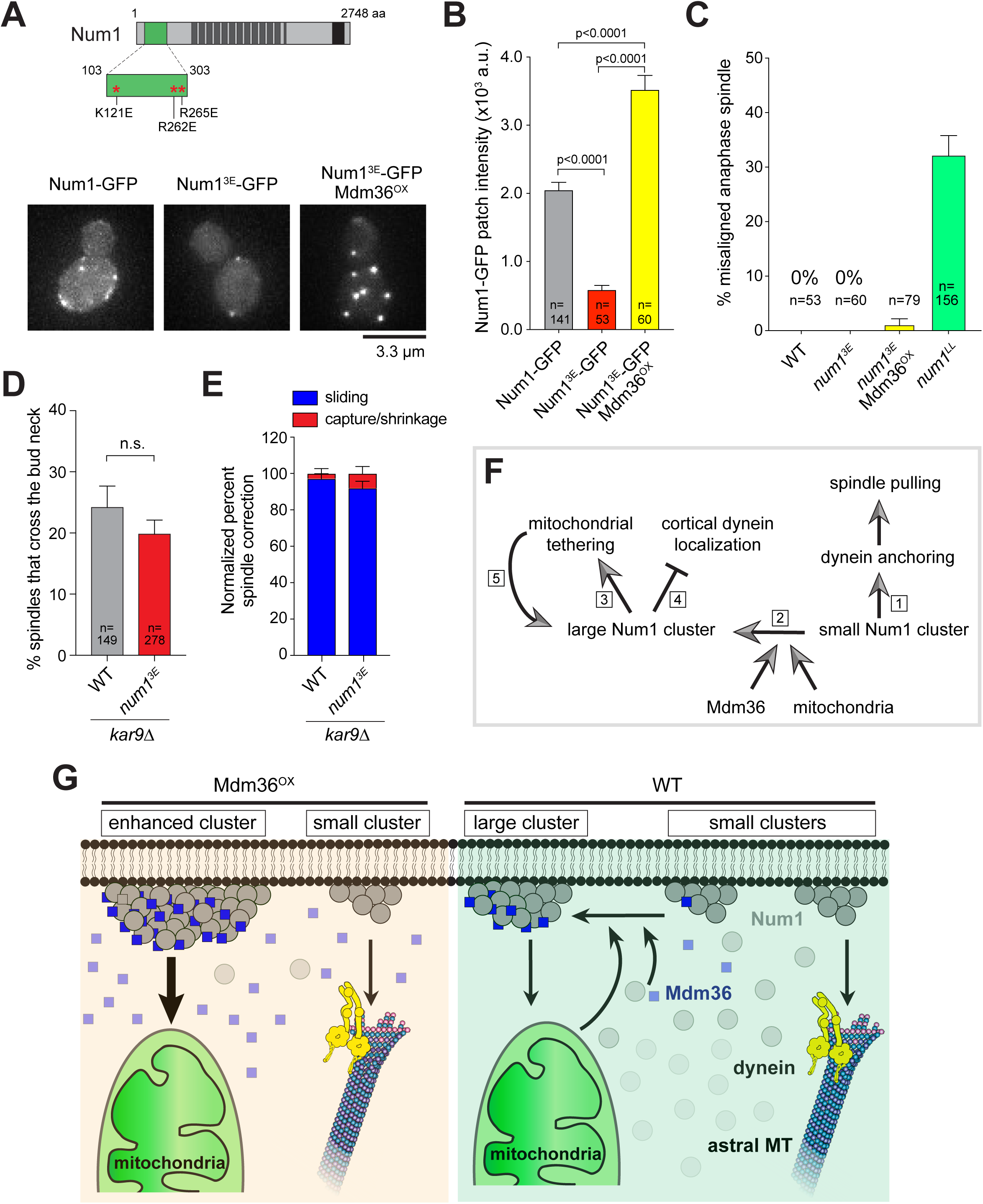
Num1^3E^ forms dynein-functioning cortical patches. (A) Schematic diagram indicating location of 3E mutations in Num1. *Bottom*, representative images of Num1-GFP and Num1^3E^-GFP in WT or Mdm36^OX^ background. Each image is a maximum intensity projection of 5 optical sections spaced 0.5 µm apart. (B) Intensity of Num1-GFP patches for strains indicated in (A). Error bar represents SEM. *P*-value by unpaired *t* test. (C) Percentage of anaphase spindles with a misoriented phenotype for indicated strains (53 ≤ n ≤ 418 spindles per strain). Error bar represents SEP. (D) Percentage of HU-arrested spindles that crossed the bud neck during a 10-min movie. Error bar represents SEP (149 ≤ n ≤ 278 spindle per strain). n.s., not statistically significant by unpaired *t* test. (E) Quantification of spindle correction mechanisms (36 ≤ n ≤ 50 events per strain). Error bar represents SEP. (F) Differential functions of small and large Num1 clusters. Small Num1 clusters mediate dynein anchoring and cortical spindle pulling function (1). Mdm36 and mitochondria cooperatively mediate formation of large Num1 clusters (2) for mitochondrial tethering function (3). Enhancing Num1 patches reduces cortical dynein targeting (4), probably because Mdm36 and mitochondria both bind to the same CC domain of Num1 as dynein. As proposed (Kraft and Lackner, 2017), mitochondria in turn promotes large cluster formation and stabilization (5). (G) Model for Mdm36/mitochondria-dependent enhancement of Num1 clustering. Diagram illustrates scenarios in WT and Mdm36OX cells. In both cases, two morphologically and functionally distinct populations of Num1 clusters (i.e. small and large clusters) mediate spindle pulling and mitochondrial tethering functions at the cell cortex. See (F) and text for further explanations.

We introduced the 3E mutations into full-length GFP-tagged Num1 and expressed the fusion protein from its endogenous locus in the WT and Mdm36^OX^ backgrounds. WT cells expressing Num1^3E^-GFP displayed fewer (Fig. S2B) and dimmer cortical patches (Fig. 5A and B) compared with those expressing Num1-GFP. However, similar to WT Num1, the intensity of Num1^3E^-GFP patches was dramatically enhanced upon Mdm36 overexpression (Fig. 5A and B), consistent with the notion that the 3E mutations did not disrupt the interaction of Num1 with Mdm36 (Ping et al., 2016). Surprisingly, spindle orientation analysis revealed that dynein pathway function was unaffected in Num1^3E^-GFP cells (0% versus 0% and 32.1% of misaligned anaphase spindle phenotype for *num1*^*3E*^ versus WT and the control strain *num1*^*LL*^, respectively; Fig. 5C) regardless of Mdm36 overexpression (compare *num1*^*3E*^ with *num1*^*3E*^ Mdm36^OX^; Fig. 5C), supporting the idea that small and dim Num1^3E^ foci are sufficient for dynein-based spindle-pulling activity. Moreover, tetrad dissection analysis showed that *num1*^*3E*^ *kar9*Δ progeny formed viable haploid colonies (Table I), whereas *num1*^*LL*^ *kar9*Δ progeny were inviable, indicating that small and dim Num1^3E^ foci are sufficient for rescuing the synthetic lethality with *kar9*Δ. Thus, our genetic and cell biological data argue that although Num1-mitochondria interaction is required for Num1 clustering (namely the formation of bright Num1 patches), the assembled patches are dispensable for dynein pathway function.

To interrogate dynein function in *num1*^*3E*^ cells more closely, we monitored preanaphase spindle movements through the bud neck using a spindle crossing assay (Fig. 5D) and assessed dynein-dependent astral MT interaction with the bud cortex using a spindle correction assay (Fig. 5E). Consistent with an intact dynein pathway function, the frequency of observing the preanaphase spindle traversing the bud neck in *num1*^*3E*^ *kar9*Δ cells in a spindle crossing assay was indistinguishable from the control *kar9*Δ cells (19.8% vs. 24.2% for *num1*^*3E*^ *kar9*Δ and *kar9*Δ, respectively, p = 0.287; Fig. 5D). Additionally, as observed in *kar9*Δ cells, the re-alignment of anaphase spindles from a misoriented position in *num1*^*3E*^ *kar9*Δ cells was predominantly mediated by MT sliding along the bud cortex (92.0% vs. 97.2% for *num1*^*3E*^ *kar9*Δ and *kar9*Δ, respectively, p = 0.3080; Fig. 5E), further supporting the notion that Num1 does not need to assemble into a large focal patch to enable cortical force generation by dynein. Consistent with this idea, two-color time-lapse movies of spindle correction showed that MT sliding occurred upon interaction of the plus end with a dim Num1^3E^-GFP focus at the bud cortex (Video 6). Furthermore, co-imaging with Cse4-GFP cells revealed that the median protein copy number for Num1^3E^-GFP foci mediating spindle correction in Mdm36^OX^ *kar9*Δ cells was 5.6 (Fig. S2C), a number strikingly similar to that observed for WT Num1-GFP foci (Fig. 4C). These results show that a single mitochondria-independent Num1 patch containing a few Num1 molecules is sufficient for dynein-based spindle pulling activity at the bud cortex.

We next investigated whether Num1’s mitochondrial tethering function is affected by the 3E mutations in Mdm36^OX^ cells. The majority of *num1*^*3E*^ cells displayed a collapsed mitochondrial network (Fig. S2A) (Ping et al., 2016), in contrast to the reticulated and cortically-tethered network observed in WT cells. Remarkably, the majority of *num1*^*3E*^ Mdm36^OX^ cells (containing enhanced Num1^3E^ patches; see Fig. 5A) displayed a WT mitochondrial network (Fig. S2A). The percentage of *num1*^*3E*^ Mdm36^OX^ cells showing a collapsed mitochondrial phenotype was significantly less than that exhibited by *num1*^*3E*^ cells (5.3% versus 78.0% for *num1*^*3E*^ Mdm36^OX^ and *num1*^*3E*^, respectively), indicating that mitochondrial tethering function was restored by enhancing Num1^3E^-GFP patches. This result indicates that Mdm36-mediated Num1 clustering is critical for Num1’s function in mitochondrial tethering.

## DISCUSSION

Here we provide evidence demonstrating that Mdm36 levels determine the size of Num1 clusters at the cell cortex. We speculate that proper clustering of Num1 is mediated through cooperative actions of Mdm36 and mitochondria (Fig. 5F and G), since both *mdm36*Δ deletion (Fig. 1A and S2D) and 3E mutations (Fig. 5A and S2D) resulted in a Num1 clustering phenotype, and that the phenotype of 3E mutations was rescued by Mdm36 overexpression (Fig. 5A and B; Fig. S2A and D). Importantly, the current view on Num1 assembly (Kraft and Lackner, 2017; Ping et al., 2016; Schmit et al., 2018) postulates that cortical clustering of Num1 serves to simultaneously anchor mitochondria and dynein to the plasma membrane. Our results, however, show that enhancing Num1 clustering only intensifies mitochondrial tethering function of Num1, not dynein-based spindle pulling function, which is inconsistent with a simultaneous interaction with dynein and mitochondria. This inconsistency is underscored by (1) the lack of strong colocalization between cortical Dyn1-3GFP and the enhanced Num1 patches seen tethering mitochondria in Mdm36^OX^ cells, (2) the fact that astral MT sliding was rarely observed to occur on a brightly enhanced Num1 patch, and (3) the lack of spindle misorientation phenotype in *num1*^*3E*^ cells where mitochondria-Num1 interaction was disrupted.

One possible explanation is that dynein anchorage and cortical force generation might be performed by a distinct population of dim Num1 clusters unaffected by Mdm36 overexpression, an idea consistent with the lack of correlation between patch enhancement and hyperactivation of cortical dynein activity (Fig. 4E). The existence of this population of Num1 is supported by direct observation of individual encounters between a dynamic astral MT plus end and a dim Num1 cluster at the bud cortex (Fig. 4B) and with the ensuing MT sliding occurring without Mdm36 being present at the cluster despite the fact that Mdm36 was overexpressed in the cell. We do not think that these observations are caused by artificial overexpression of Mdm36, since we also observed MT sliding occurring in *mdm36*Δ cells, where Num1 foci are similarly small in size. We speculate that the redistribution of Num1 molecules observed in Mdm36^OX^ cells, as compared to WT cells (Fig. 2A), makes it possible to follow astral MT interaction with individual cortical Num1 sites. Such encounters would have been otherwise impossible to observe in WT cells due to the abundance and heterogeneity of Num1 clusters along the cell cortex (see Figure 1A in Omer et al., 2018, Figure 3A in Heil-Chapdelaine et al., 2000, Figure 1A in Kraft and Lackner, 2017, and Figure 1C in Schmit et al., 2018).

The budding yeast model is uniquely suited for *in vivo* assessment of spindle pulling under load. The movement of the spindle into the narrow aperture of the bud neck introduces a load burden that antagonizes dynein pulling at the bud cortex. How much force is needed to pull the spindle across the bud neck remains a total mystery. Our molecule-counting data (Fig. 4C and Fig. S2C) suggest that a small ensemble of approximately six Num1 molecules is sufficient for dynein anchorage and cortical force generation under load. Assuming Num1 binds dynein with a 1:1 stoichiometry, an ensemble of six Num1 molecules (which probably exist as three Num1 dimers (Ping et al., 2016; Tang et al., 2012)) can potentially anchor up to three dynein dimers. Since each dynein dimer produces 4-7 pN of force (Belyy et al., 2016; Cho et al., 2008; Gennerich et al., 2007), an ensemble of Num1 anchoring three dynein dimers can generate a force of 3 x 4-7 = 12-21 pN, which we speculate likely corresponds to the minimum force needed to pull the spindle across the bud neck. By comparison, in *Dictyostelium* cells, 8-16 pN of dynein-generated force is needed for rapid transport of maturing phagosomes to lysosomes (Rai et al., 2016). In these cells, the phagosomes acquire increasingly greater minus end-directed movements (as they mature) through clustering of multiple dynein motors (up to 16) into a microdomain on the phagosomal surface (Rai et al., 2016). Thus, unlike the evidence for *Dictyostelium* dynein in phagosomal transport, our data suggest that clustering of a large number of motors into a single site does not appear to be a key regulatory step for yeast dynein’s function in spindle positioning. In cultured human cells, although cortical clustering behavior has recently been demonstrated for the dynein-anchoring homologue NuMA (Okumura et al., 2018), the size of the NuMA clusters and the force required for spindle displacement remain to be determined. In future studies, it would be interesting to test whether small NuMA clusters can generate large spindle-pulling forces, as observed in yeast.

## MATERIALS AND METHODS

### Media and Strain Construction

Yeast media was obtained from Sunrise Science Products (San Diego, CA). All strains (Table S1) were generated in the background of YWL36 (Vorvis et al., 2008) by standard genetic crosses or PCR product-mediated homologous recombination at the chromosomal locus (Longtine et al., 1998). Yeast was transformed using the lithium acetate method (Knop et al., 1999). To overexpress Mdm36, we replaced 50 nt of *MDM36* promoter immediately upstream of the start codon with a hygromycin resistance marker (HPH) and 562 bp of *MET3* promoter using the plasmid pHPH::MET3p as PCR template. To generate Num1^3E^ (K121E, R262E and R265E) expressed from the endogenous chromosomal locus, we used the site-specific genomic mutagenesis approach (Gray et al., 2004). The CC domain (amino acid 25-290) was first replaced in a Num1-GFP strain with *URA3* marker from pRS306. Next, we generated a PCR fragment containing the mutated residues using an overlap extension PCR method. We included flanking sequences for homologous recombination targeting the corresponding chromosomal region that has been replaced by the *URA3* marker. Transformants were plated on media containing 5-fluoroorotic acid (5-FOA) to select for substitution of *URA3* with the PCR fragment. We then verified all point mutations by DNA sequencing of the genomic locus. To overexpress mRuby2-Mdm36 (tagged at its N-terminus) using the *MET3* promoter, a 1347-bp 5’ fragment of the *MDM36* open reading frame was subcloned into a *LEU2* plasmid (pBJ78) containing *MET3p*:mRuby2. The resulting plasmid was linearized by BamHI (within the *MDM36* open reading frame) and transformed into a haploid strain to integrate at the endogenous *MDM36* locus. To label MTs, strains were transformed with Tub1-tagging plasmids for integration into *TUB1* locus as previously described (Markus et al., 2015).

### Image Acquisition and Analysis

Wide-field fluorescence images were acquired at room temperature using a 1.49 NA 100x objective on a Nikon TiE inverted microscope equipped with a Nikon LUN4 laser unit (405 nm, 488 nm, 561 nm, and 640 nm), an EMCCD camera (iXon 888; Andor), and a sCMOS camera (Zyla 4.2; Andor). We used a multi-pass quad filter cube set (C-TIRF; Chroma) for imaging GFP, Venus, and mRuby2/mRuby3/mCherry fluorescence. Microscope system was controlled by NIS-Elements imaging software (Nikon). Image stacks were deconvolved where indicated using 3D Deconvolution in NIS-Elements with the Automatic method. We performed standard photobleaching correction where indicated using Bleach Correction in Fiji with the Simple Ratio method. Cells were grown to mid-log phase in nonfluorescent synthetic defined (SD) media at 30°C and mounted on a 1.7% agarose pad containing SD or SD lacking methionine for imaging. To induce the *MET3* promoter, overnight cultures grown at 30°C in SD media were harvested, washed once with water, diluted into fresh SD media lacking methionine, and grown for 5-6 hours at 30°C before imaging. Confocal images were acquired using a 1.4 NA 60X oil immersion objective on an Andor W1 Spinning Disk Confocal microscope housed in the Life Sciences Center Imaging Facility at Dartmouth College.

To determine the copy number of Num1-GFP at individual cortical patches, we used the ratiometric comparison of fluorescence intensity approach as described in Verdaasdonk et al. (2014). Strains expressing Num1-GFP were mixed with the Cse4-GFP strain and imaged simultaneously in the same field. We used Fiji to draw a square encompassing 4 x 4 pixels (Fig. 1C) or 3 x 3 pixels (Fig. 4C and Fig. S2C) to quantify the integrated intensity of individual anaphase Cse4-GFP clusters and a 3 x 3 pixel square for quantifying cortical Num1-GFP clusters. To subtract the background intensity from each measurement, we moved the square from the cluster to a nearby cytoplasmic area within the same cell. The mean integrated intensity of Cse4-GFP clusters was assigned a value of 96 molecules (Lawrimore et al., 2011; Verdaasdonk et al., 2014) and was used to calculate the copy number of Num1-GFP.

To measure the number of bright cortical Num1 patches per cell, we performed particle analysis using the Analyze Particles tool in Fiji with the minimum particle size set at 9 pixel units and the threshold set at the background intensity of the cytoplasm. To measure the total fluorescence intensity of Dyn1-3GFP and Jnm1-3mCherry at the plus end and cell cortex, we used the circle selection tool in Fiji to encompass the 3GFP or 3mCherry spot and measured the intensity from the maximum intensity projection images. Background intensity from a nearby cytoplasmic area was subtract from each measurement.

To measure spindle velocity, we used the mTrack plugin in Fiji to track the position of the SPB (determined from the mRuby2-Tub1 image) during a 7-min movie in cells arrested with hydroxyurea (HU). We grew the cells to mid-log phase and arrested them with 200 mM HU for 1-1.5 h before imaging. For spindle penetration assay, we measured the farthest distance traveled by a spindle pole moving across the bud neck during a 10-min movie.

To plot the intensity profile of Num1-GFP along the cell cortex, we used the segmented line tool in Fiji to trace the bud and mother cell perimeter. Next, we used the Plot Profile tool to graph the intensity along the segmented line.

For spindle correction assay, we scored for misaligned anaphase spindles that moved into the bud neck and became aligned along the mother-bud axis during a 10-min movie in *kar9*Δ background. Spindle correction was scored as mediated by “sliding mechanism” or “capture-shrinkage mechanism” based on the interaction displayed by the astral MT with the bud cortex while the spindle moved into the bud neck during its realignment, as previously described (Omer et al., 2018).

### Sucrose Gradient, Cell Lysis, and Western Blotting

Sedimentation analysis on sucrose gradients was performed as previously described (Omer et al., 2018). For analysis of Num1-GFP levels, overnight yeast cultures grown at 30°C in 15 ml SD media lacking methionine were harvested and resuspended in ice-cold lysis buffer containing 20 mM Tris pH 7.5, 150 mM NaCl, 1 mM EDTA, and 1.5% Triton X-100 supplemented with protease inhibitor tablet (Millipore Sigma). Equal amounts of cells were lysed by bead beating six times for 30 s, with 2 min on ice between each beating. Next, we centrifuged the crude lysate at 500 *g* for 5 min at 4°C in a microfuge (Eppendorf, model 5424R) to remove cell debris. The resulting supernatants were separated on 4-15% SDS-PAGE gels (Bio-Rad) and then electro-blotted to nitrocellulose membrane using a Trans-Blot Turbo Transfer System (Bio-Rad). Membranes were probed with rabbit anti-GFP antibody (Chromotek) at 1:1000, rabbit anti-Sac1 antibody (a gift from Dr. Charles Barlowe) at 1:2000, and HRP-conjugated goat anti-rabbit antibody (Jackson ImmunoResearch) at 1:10,000 dilution. Chemiluminescence signals were acquired using a ChemiDoc Imaging System (Bio-Rad). Immunoblots were exposed for durations ranging from 1 s to 10 min without saturating the camera’s pixels.

### Statistical Methods

All statistical significance in this study was determined by unpaired Student’s *t* test or one-way ANOVA test performed using the GraphPad Prism software.

## Supporting information

Omer et al_video 1

Omer et al_video 2

Omer et al_video 3

Omer et al_video 4

Omer et al_video 5

Omer et al_video 6

## ACKNOWLEDGEMENT

This work was supported by an NIH/NIGMS grant (GM076094) to W.L. Lee and in part by a Scholarly Innovation and Advancement Award at Dartmouth College to W.L. Lee. We thank Samuel Greenberg and Weimin Tan for valuable help with making yeast strains.

The authors declare no competing financial interests.

## SUPPLEMENTARY LEGENDS

**Table S1.**
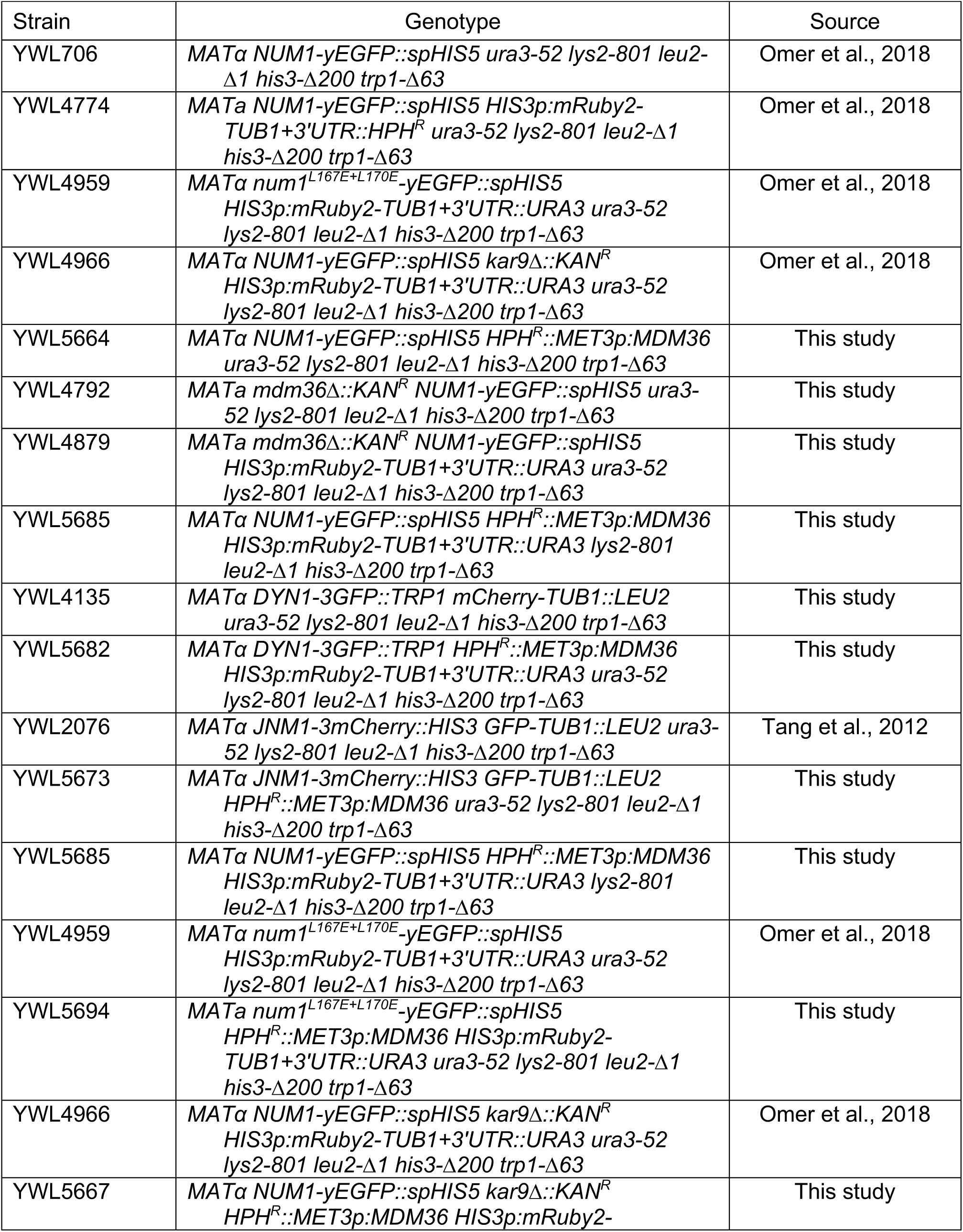
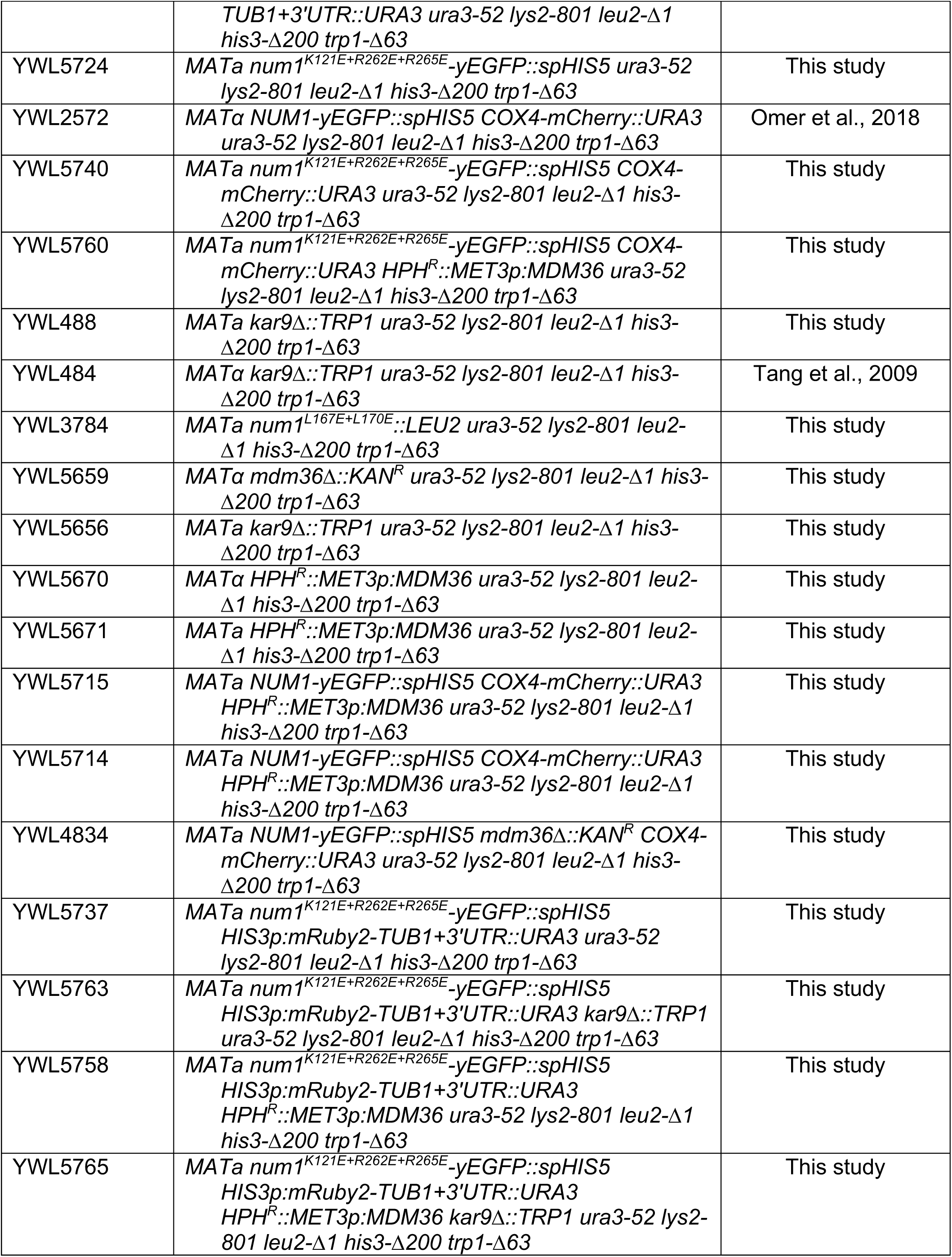
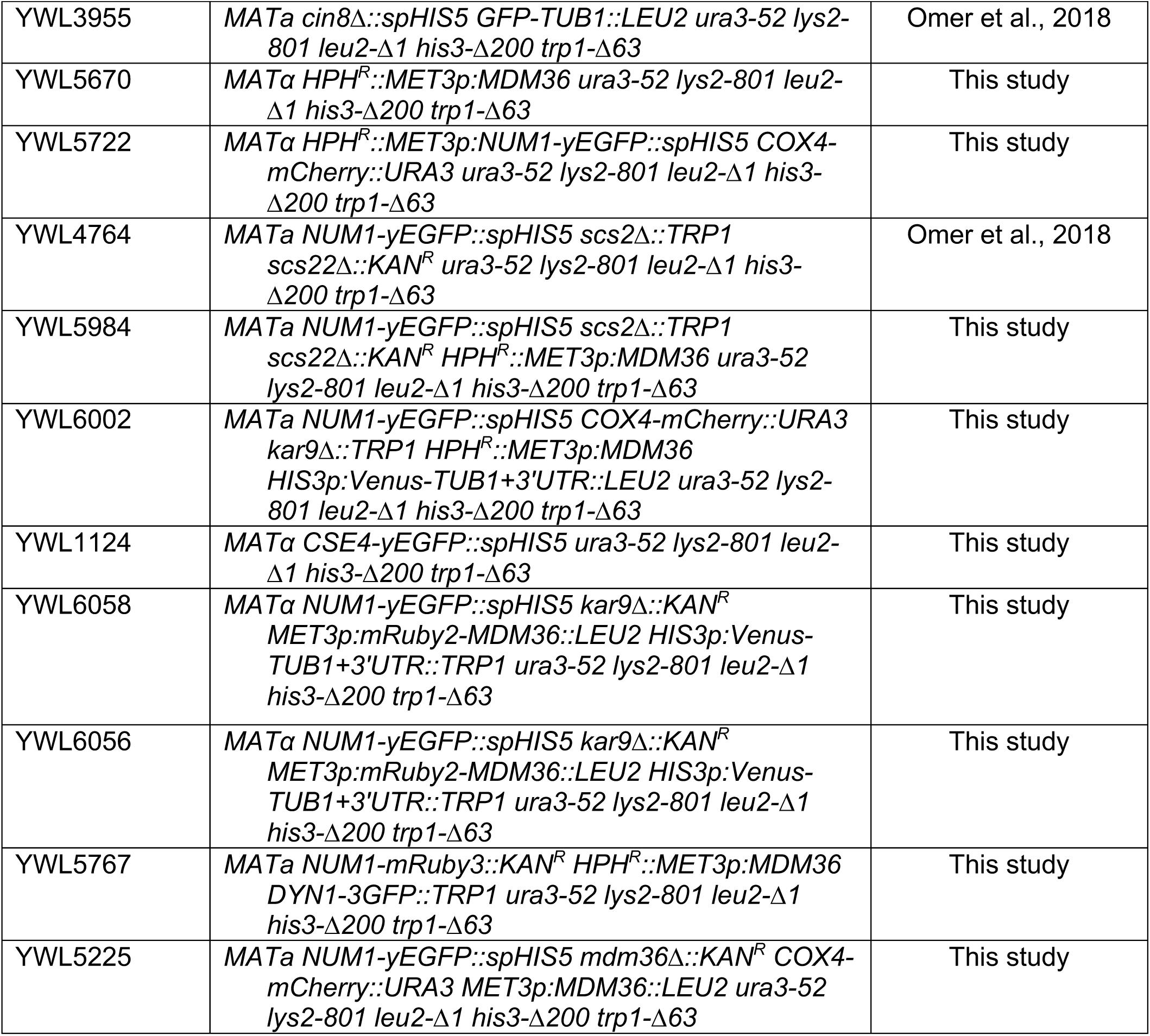
Yeast strains used in this study

**Figure S1.**
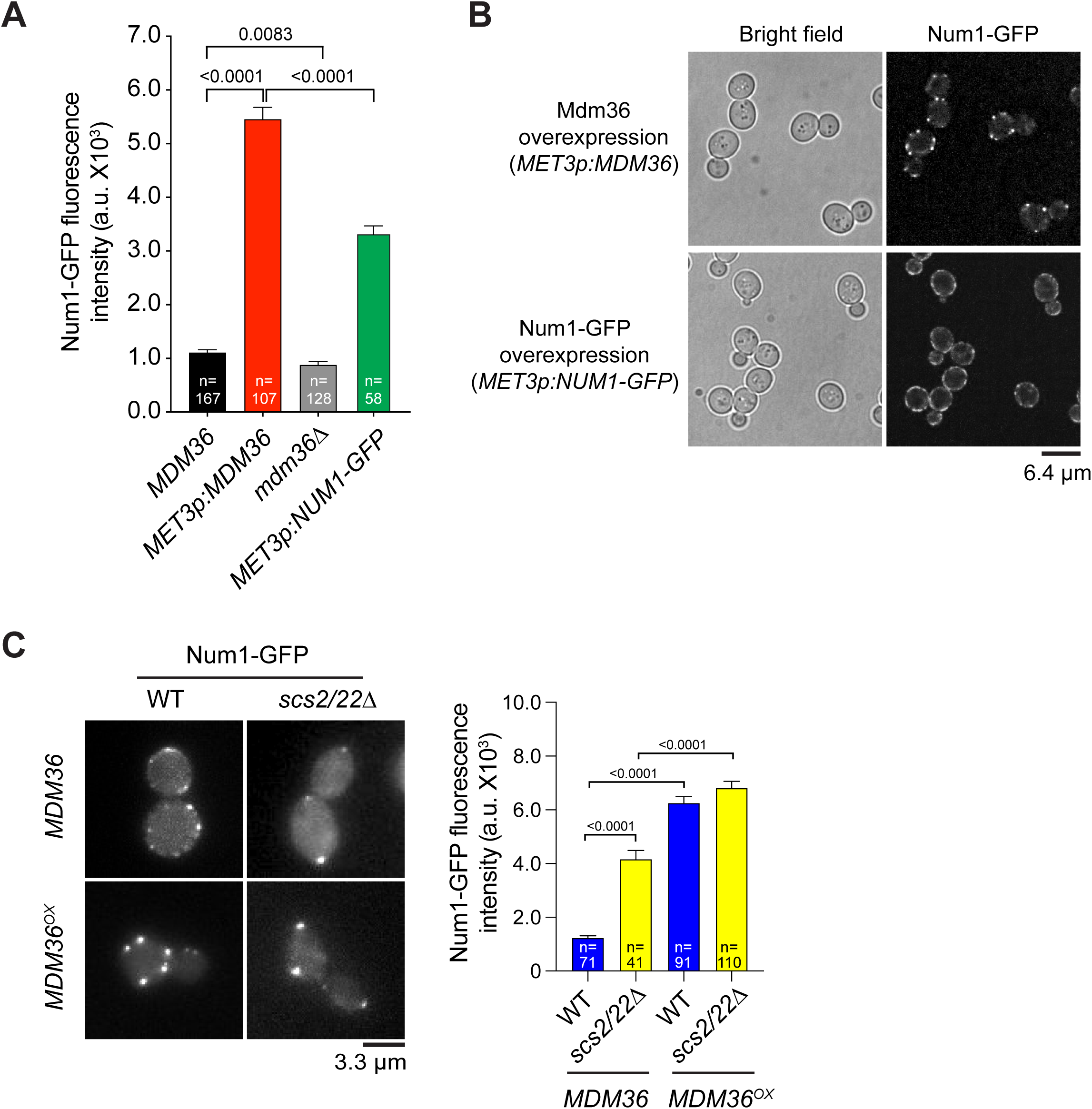
Mdm36 promotes Num1 clustering. (A) Mean fluorescence intensity of individual Num1-GFP patches in indicated strains. Error bars represent SEM. P-value by unpaired *t* test. (B) Bright field and confocal images of cells overexpressing Mdm36 or Num1-GFP. Each confocal image is a maximum intensity projection of four optical sections spaced 0.2 µm apart from the equator of cells. **Num1 redistributes to bud tip and mother apex in Mdm36**^**OX**^ **cells lacking Scs2 and Scs22**. (C) Wide-field images of Num1-GFP in WT and *scs2/22*Δ cells with and without Mdm36 overexpression. Each image is a maximum intensity projection of five optical sections spaced 0.5 µm apart. Plot showing mean fluorescence intensity of individual Num1-GFP patches in indicated strains. Error bars represent SEM. P-value by one-way ANOVA test.

**Figure S2.**
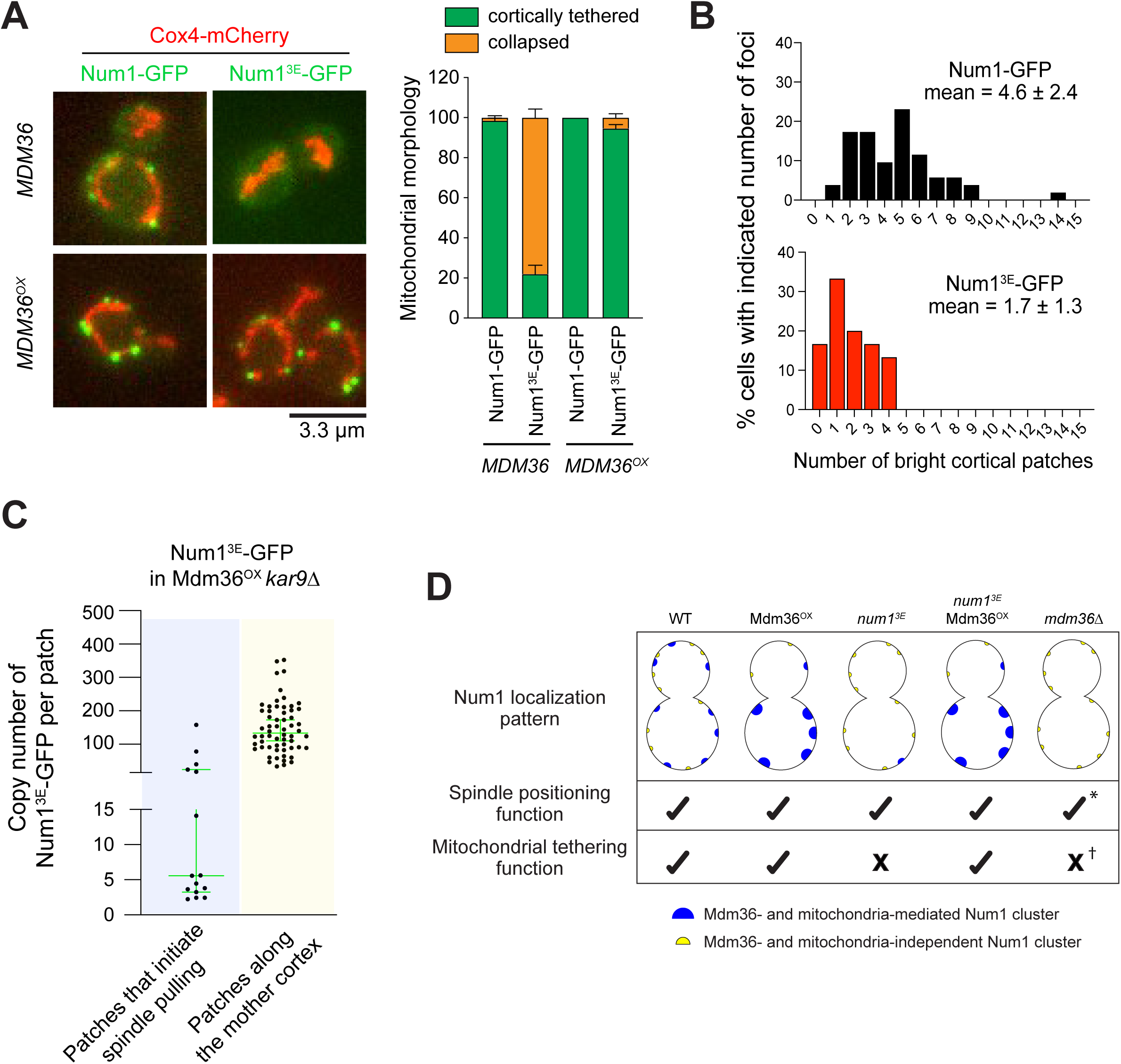
3E mutations disrupt mitochondrial tethering but not spindle positioning function of Num1. (A) Wide-field images of Cox4-mCherry cells expressing Num1-GFP and Num1^3E^-GFP with and without Mdm36 overexpression. *Right*, quantification of mitochondrial morphology in indicated strains. Mitochondrial network is considered collapsed if the network fails to show tubular morphology and appears meshed or aggregated over the course of a 4-min movie. Error bars represent SEP. 91 ≤ n ≤ 131 cells per strain. (B) Percentage of Num1-GFP and Num1^3E^-GFP cells with indicated number of cortical patches (30 ≤ n ≤ 52 cells). Mean number per cell ± SD is indicated. (C) Copy number of Num1^3E^-GFP at individual cortical patches found along the mother cell cortex compared to those found in the bud cortex mediating spindle pulling during a spindle correction assay in Mdm36^OX^ *kar9*Δ cells (n = 60 and 15 patches, respectively). Median ± 95% confidence interval is indicated. (D) Graphical summary of Num1 localization pattern and the resultant spindle positioning and mitochondrial tethering function in the indicated strain backgrounds. *, based on viability of *mdm36Δ kar9*Δ tetrad progeny (Table I). †, based on mitochondrial phenotype of *mdm36*Δ (Hammermeister et al., 2010).

## SUPPLEMENTARY VIDEO LEGENDS

**Video 1. Enhanced Num1 patches in Mdm36**^**OX**^ **cells are stationary and remarkably stable**. Time-lapse images of Num1-GFP in a Mdm36^OX^ cell showing enhanced Num1 patches persisting for greater than two cell division cycles. Each frame is a maximum intensity projection of 9 optical sections spaced 0.5 µm apart. Movie captured at 10 min intervals.

**Video 2. MT sliding initiates at a dim Num1 patch**. Time-lapse images of Num1-GFP and mRuby2-Tub1 in Mdm36^OX^ *kar9*Δ cells showing examples of MT sliding events initiated by the plus end contacting a dim Num1 patch (arrowheads) at the bud cortex, followed by spindle correction. Each frame is a maximum intensity projection of 5 wide-field sections spaced 0.5 µm apart. Movie captured at 10 s intervals.

**Video 3. MT sliding occurring at a bud cortical region without visible Num1-GFP**. Time-lapse images of Num1-GFP and mRuby2-Tub1 in Mdm36^OX^ *kar9*Δ cells. Frames from 60 s to 80 s and from 130 s to 150 s show MT sliding along the bud cortex (arrows) for the bottom and top cells, respectively. Each frame is a maximum intensity projection of 5 wide-field images spaced 0.5 µm apart. Movie captured at 10 s intervals.

**Video 4. Mdm36**^**OX**^ **cells display branched and tubular mitochondrial network**. Full 3D reconstruction of a deconvolved wide-field image stack showing a cell expressing Num1-GFP and mitochondria-targeted Cox4-mCherry in Mdm36^OX^ background. Image stack consists of 11 optical sections spaced 0.5 µm apart.

**Video 5. Mitochondrial network is tethered by enhanced Num1 foci in Mdm36**^**OX**^ **cells**. Full 3D reconstruction of a deconvolved wide-field image stack showing mitochondria association with every single enhanced Num1-GFP patches in a Mdm36^OX^ cell. Mdm36 was overexpressed using a *MET3p:MDM36* plasmid integrated at the *leu2* locus. Image stack consists of 31 optical sections spaced 0.2 µm apart encompassing the entire thickness of the cell.

**Video 6. MT sliding initiates at a dim Num1**^**3E**^ **patch**. Time-lapse images of Num1^3E^-GFP and mRuby2-Tub1 in a *kar9*Δ cell. Frames at 0 s through 80 s show a dim cortical Num1^3E^-GFP patch (arrowhead) that photobleached in later frames. Frame at 130 s shows the plus end contacting the site of the dim Num1^3E^-GFP patch, followed by MT sliding in the subsequent frame. Each frame is a maximum intensity projection of 5 wide-field sections spaced 0.5 µm apart. Movie captured at 10 s intervals.

